# Modelling the effect of ephaptic coupling on spike propagation in peripheral nerve fibres

**DOI:** 10.1101/2021.11.30.470540

**Authors:** Helmut Schmidt, Thomas R. Knösche

## Abstract

Experimental and theoretical studies have shown that ephaptic coupling leads to the synchronisation and slowing down of spikes propagating along the axons within peripheral nerve bundles. However, the main focus thus far has been on a small number of identical axons, whereas realistic peripheral nerve bundles contain numerous axons with different diameters. Here, we present a computationally efficient spike propagation model, which captures the essential features of propagating spikes and their ephaptic interaction, and facilitates the theoretical investigation of spike volleys in large, heterogeneous fibre bundles. The spike propagation model describes an action potential, or spike, by its position on the axon, and its velocity. The velocity is primarily defined by intrinsic features of the axons, such as diameter and myelination status, but it is also modulated by changes in the extracellular potential. These changes are due to transmembrane currents that occur during the generation of action potentials. The resulting change in the velocity is appropriately described by a linearised coupling function, which is calibrated with a biophysical model. We first lay out the theoretical basis to describe how the spike in an active axon changes the membrane potential of a passive axon. These insights are then incorporated into the spike propagation model, which is calibrated with a biophysically realistic model based on Hodgkin-Huxley dynamics. The fully calibrated model is then applied to fibre bundles with a large number of axons and different types of axon diameter distributions. One key insight of this study is that the heterogeneity of the axonal diameters has a dispersive effect, and that with increasing level of heterogeneity the ephaptic coupling strength has to increase to achieve full synchronisation between spikes. Another result of this study is that in the absence of full synchronisation, a subset of spikes on axons with similar diameter can form synchronised clusters. These findings may help interpret the results of noninvasive experiments on the electrophysiology of peripheral nerves.

## 1 Introduction

Signal transmission in peripheral nerve fibres is based on the propagation of action potentials, or spikes, along the axonal membrane, which alters the electrophysiological properties of the extracellular medium and therefore influences nearby axons via so-called *ephaptic coupling* [1–3]. Early experiments have demonstrated that in the presence of a highly resistive extra-cellular medium, action potentials travelling in two parallel, closely spaced axons synchronise and travel at a lower velocity than in the isolated case [4–7]. This has been reproduced in theoretical work using numerical or analytical tools [8–18]. Both theoretical and experimental work, however, has been restricted thus far to a small number of identical axons due to the experimental or computational effort (with the exception of [18]). In contrast, peripheral nerve bundles are composed of a relatively large number of nerve fibres, which follow a wide distribution of axonal diameters [19–21]. As there is an approximately linear relationship between the axon diameter and the velocity of a spike [16, 22, 23], one can expect that structural heterogeneity counteracts the synchronisation of spikes, akin to phase-coupled oscillators, where wider distributions of the angular frequencies require stronger coupling to achieve synchronisation [24, 25]. The main difference between phase-coupled oscillators and ephaptically coupled spikes is that in the latter case the coupling is restricted to axonal segments close to the spikes, and that the lifetime of a spike is restricted to the time interval between spike generation and the spike reaching the distal end of the axon. Therefore, spike synchronisation depends crucially on the initial conditions, as demonstrated recently for white-matter fibre bundles [26]. For this reason we focus our analysis on initially synchronous spike volleys, in line with previous studies [12, 13].

Our aim is to systematically study ephaptic coupling effects in peripheral nerves with large axon counts and distributed morphology. For this, we devise a simplified spike-propagation model (SPM) that reduces the computational effort significantly as compared to solving models based on nonlinear partial differential equations (PDEs). The SPM is inspired by phase-coupled oscillators in the sense that each spike is endowed with an intrinsic propagation velocity based on the morphology of the associated axon. Furthermore, each spike can be ascribed a coupling function that characterises the spatio-temporal profile of ephaptic coupling with spikes in nearby axons. The collective ephaptic coupling generated by all spikes in a fibre bundle then perturbs the intrinsic velocity of a single spike, which decelerates if the axonal membrane is hyperpolarised, and vice versa. In the resulting numerical scheme, the position of each spike is propagated based on its perturbed velocity. In turn, the positions and velocities of the spikes determine their mutual ephaptic coupling effects.

We begin our investigation by considering an exemplary bundle containing two axons, for which we develop the theoretical basis that can be extended to larger fibre bundles. First, we compute the effect that the spike in an active axon has on a nearby passive axon, which we do analytically under the assumption that the axons are homogeneous and that the perturbation of the passive axon can be captured by the linear terms of the cable equation. As most axons in peripheral nerves are myelinated, the assumption of homogeneity implies that the specific location of nodes of Ranvier is irrelevant. We calibrate the proposed spike propagation model using a biophysically realistic model based on Hodgkin-Huxley dynamics.

The calibrated SPM is then used to study large axon bundles systematically, especially the interplay between axon diameter distribution and the strength of ephaptic coupling, with the latter having a synchronising effect, and the former a dispersive effect. The coupling strength is determined by the fibre density (i.e. relative volume occupied by nerve fibres), and the resistivity of the extracellular medium. The aim here is to identify conditions under which synchronisation can occur given different axon diameter distributions and structural parameters, especially fibre density.

## 2 Ephaptic coupling between two nerve fibres

In this section we lay the theoretical basis of the SPM by considering a bundle of two nerve fibres. First we derive an analytical expression for the extracellular potential (or local field potential) generated by a spike. This is then used for the perturbation of the membrane potential of a passive axon generated by a spike in a nearby axon, which can be regarded as the ephaptic coupling between two nerve fibres. Next, we introduce a piecewise quadratic approximation of the spatial spike profile, which allows us to compute the ephaptic coupling function analytically. Lastly, we calibrate the SPM by fitting free parameters to a biophysically realistic model.

### 2.1 Perturbation of the extracellular medium by a spike in a single axon

Axons may be regarded as core conductors [27, 28], which allows to describe the formation and propagation of spikes by the one-dimensional cable equation:

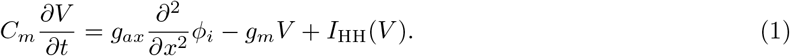

The derivation of the cable equation is based on Kirchhoff’s first law, which states that the currents are balanced at any given location inside or outside the axon. Here, the membrane potential *V* is defined as the difference between intracellular and extracellular potential, *V* = *ϕ*_*i*_ − *ϕ*_*e*_. If the extracellular medium is grounded, then *V* = *ϕ*_*i*_ and the traditional cable equation is recovered. In general, however, that assumption does not hold because of the finite size of the extracellular medium, and its non-zero conductivity. The term on the left describes the capacitive currents across the axonal membrane and the myelin sheath, with *C*_*m*_ being the capacitance of the membrane and myelin. The terms on the right describe resistive currents, of which the first term is the longitudinal intra-axonal resistance, which depends on the intracellular conductivity *σ*_*ax*_ and the cross-sectional area 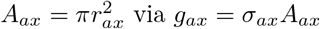. The second term is the passive resistive current across the membrane and myelin, with *g*_*m*_ being the radial resistance of the myelin and the membrane, and the third term represents voltage-gated currents described by the Hodgkin-Huxley model.

The current balance outside an axon can be formulated in the equivalent manner:

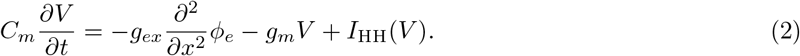

Here, the (longitudinal) extracellular resistance *g*_*ex*_ depends on the extracellular conductivity *σ*_*ex*_ and the cross-sectional area of the extracellular space, *A*_*ex*_. This relationship, however, is no longer applicable if multiple axons are in the same bundle. In this case, the extracellular potential depends on the capacitive and resistive currents generated by all axons, and we therefore find the following relationship for the current balance in the extracellular medium:

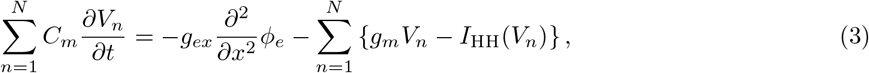

with *N* being the total number of axons. It is easy to see from Eq. (3) that if the extracellular conductivity or space is infinite, then *ϕ*_*e*_ = 0.

Typically, spikes are measured as the spatiotemporal profile of the depolarisation of the membrane potential *V*. Here, we are interested in recovering a relationship between the extracellular potential *ϕ*_*e*_ and the membrane potential *V*. Combining Eq. (3) and Eq. (1), and using *ϕ*_*i,n*_ = *V*_*n*_ + *ϕ*_*e*_, we obtain

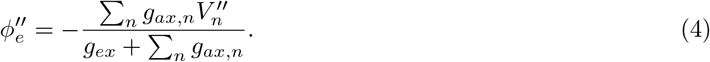

The terms 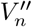 and 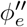 denote the second spatial derivative in the axial direction of *V*_*n*_ and *ϕ*_*e*_. Although Eq. (4) is only an implicit representation of the extracellular potential via its second spatial derivative, we show that this expression is sufficient to compute the effect of a spike onto the membrane of a nearby axon. It is also obvious that the effect of axonal activity onto the extracellular medium is additive. In the following, we use the equivalent representation in terms of conductivities and fibre density. The fibre density *ρ* can be defined as *ρ* = *g*^−2^*A*_*ax*_*/*(*g*^−2^*A*_*ax*_+ *A*_*ex*_), where *g* is the g-ratio of the fibres, i.e. axonal diameter divided by fibre diameter (axon plus myelin). The perturbation of the extracellular medium by a spike in the *n*^th^ axon is thus given by

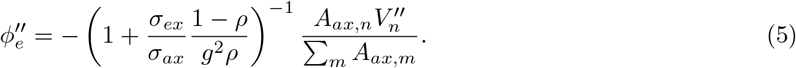

We note here that we only consider axons with circular shape, therefore *A*_*ax*_ can be replaced by the square of the fibre diameter.

### 2.2 Perturbation of the membrane potential of a passive axon by a spike in a contiguous one

A spike can be regarded as a travelling wave along the axon. If the axon is sufficiently homogeneous, then this wave appears stationary in the co-moving frame. For simplicity, we assume here that the axons under consideration are either homogeneous, which would be the case for unmyelinated axons, or homogenised in the case of periodically myelinated axons. The term homogenised refers to the technique proposed by Basser [29], which yields compound variables for the cable parameters that depend on the properties of myelinated and unmyelinated segments.

We transform the cable equation, Eq. (1), into the co-moving frame by setting *ξ* = *x* −*ct*, where *c* is the propagation velocity of the spike, which results in

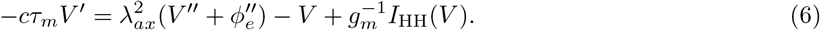

Here, we have made use again of *ϕ*_*i*_ = *V* + *ϕ*_*e*_, as well as *τ*_*m*_ = *C*_*m*_*/g*_*m*_ and 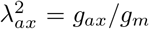, and · ′ indicates differentiation with respect to *ξ*. More specifically, *τ*_*m*_ is the homogenised time constant, and *λ*_*ax*_ is the homogenised length constant [29]:

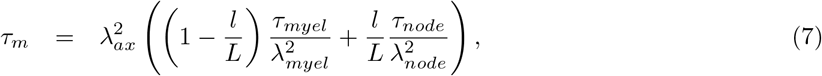

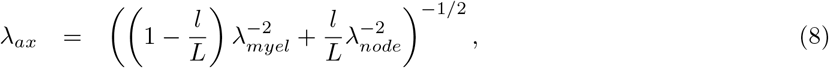

where indices denote either myelinated segments with length *L*, or nodes of Ranvier with length *l*. The node and myelin-specific parameters were chosen to be *τ*_*myel*_ = 0.47ms, *τ*_*node*_ = 0.03ms, 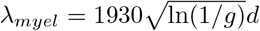, and 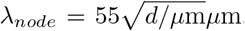, with *d* being the axon diameter. For simplicity, we set *l/L* = 0.01. Since the g-ratio is also held fixed at *g* = 0.6, the only free parameter is the axon diameter *d*.

If the perturbation is sufficiently small, it will fail to elicit a spike in the passive axon, and the nonlinear term *I*_HH_(*V*) can be neglected for simplicity. Thus we arrive at a linear ODE that describes the spatial profile of the perturbation in the passive axon:

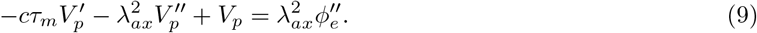

Here, *V*_*p*_ indicates the perturbation of the membrane potential caused by 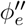. This is an inhomogeneous differential equation of second order. The homogeneous solution to (9) is found to be

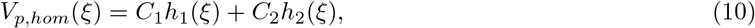

where *h*_1_(*ξ*) = exp(*ξ*/*ξ*_1_) and *h*_2_(*ξ*) = exp(*ξ*/*ξ*_2_) are the fundamental solutions to (9), with

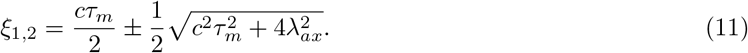

Since the perturbations are localised, we require that *V*_*p*_ → 0 as *ξ*→ ±∞, and therefore *V*_*p,hom*_ = 0. Nevertheless, the fundamental solutions serve to identify the particular solution to (9), which is found by the method of variation of parameters. The particular solution can be posed as

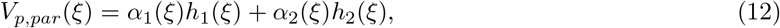

which is identical to the full solution to (9). Since differentiation will yield only one equation for the two unknowns *α*_1_(*ξ*) and *α*_2_(*ξ*), we may make a further assumption regarding the solution structure:

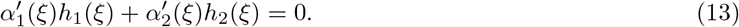

Differentiation of (12) and insertion of the resulting terms into (9) then yields

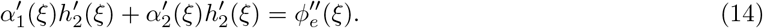

Equations (13) and (14) pose a set of two linear equations that can be solved for 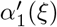 and 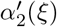. Subsequent integration results in

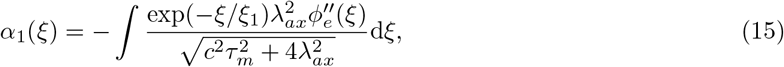

and

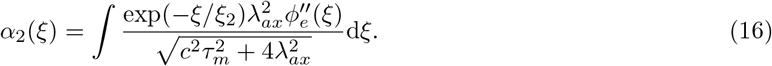

Since the integrands vanish at ±∞, we can pose these indefinite integrals as definite integrals on the interval [*ξ*, ∞) for *α*_1_(*ξ*), and (−∞, *ξ*] for *α*_2_(*ξ*) instead. This yields the following integral form for *V*_*p*_(*ξ*):

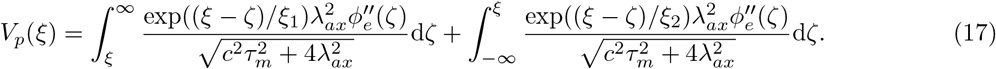

An alternative representation is the following convolution integral:

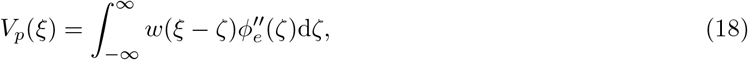

with the asymmetric convolution kernel given by

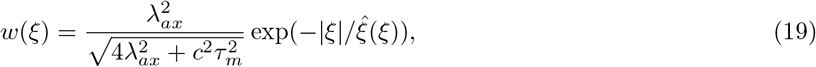

where

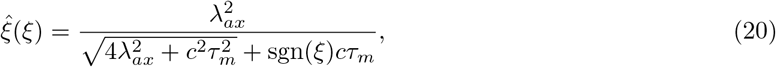

with sgn(·) being the sign function.

It is important to note here that *λ*_*ax*_ and *τ*_*m*_ are parameters of the passive axon, whereas *c* represents the velocity of the spike in the active axon. In the following we extend the notation by introducing indices for the axons that interact. As the interaction is pairwise, this notation requires two indices, one tagging the passive axon and the second one tagging the active axon. In the following example the axon with index 1 is the passive axon, and the axon with index 2 is the active axon:

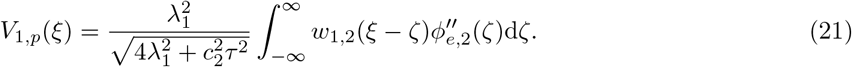

The convolution kernel and the spatial scales now also require indices:

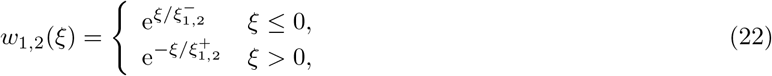

with

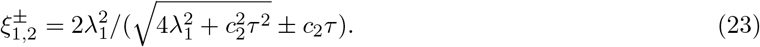

Spike profiles and the resulting perturbations can be generated numerically, upon which Eq. (21) has to be evaluated numerically as well. In Figure 1 we visualise the convolution kernel, and compare the perturbation of a passive axon that was obtained numerically, with the convolution of this kernel with the second derivative (curvature) of the spatial spike profile. The relatively small difference between the numerical and analytical result can be explained by nonlinear effects not taken into account in the convolution, and the homogenisation of the axonal parameters.

**Figure 1:**
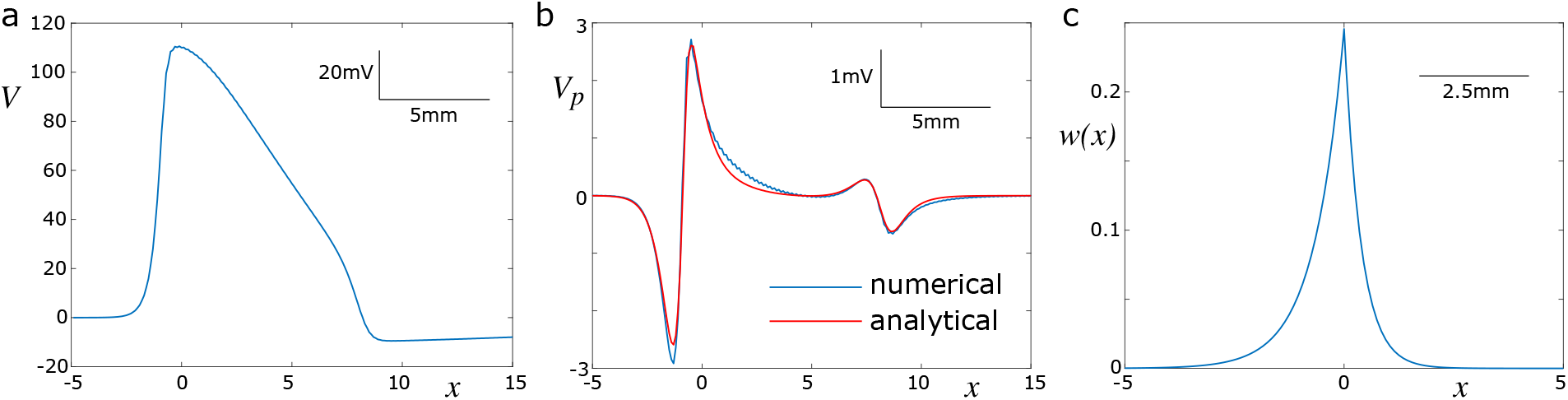
The perturbation in a passive axon can be described by a convolution of the curvature of the spike profile and an asymmetric kernel. (a) Spike profile computed numerically using the biophysical model. (b) Comparison of the perturbation in a passive axon obtained numerically with the biophysical model, and the convolution of the analytically derived integral kernel with the second derivative of the numerically obtained spike profile. (c) Profile of the asymmetric kernel. The kernel is compressed to the right (direction of propagation of the spike), and elongated to the left. Parameters: *ρ* = 0.3, *d* = 1*µ*m, *c* = 3.1m/s.

### 2.3 Piecewise quadratic approximation of spike profiles

To reduce the computational load, we opt for an approximation of the spike profile that can be adjusted via a shape parameter *a*_1_. In the time domain, the spike is parameterised as follows:

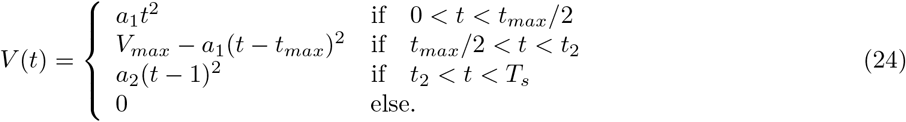

The amplitude is set to *V*_*max*_ = 110, and the other parameters can be related to *a*_1_ by imposing smoothness conditions, which results in

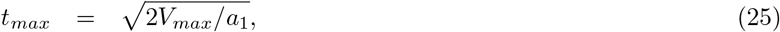

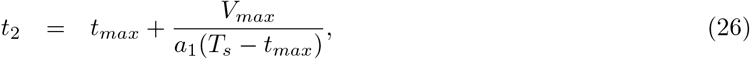

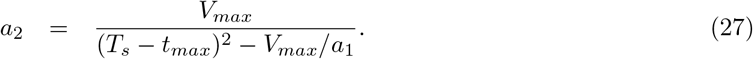

Thus, the spike has a total duration of *T*_*s*_, and its shape can be varied between a triangular shape at large values of *a*_1_, and a bell-shape at small values of *a*_1_. The shape of the spike can then be translated into the spatial domain by calling *V* (*t*) = *V* (*ξ*/*c*), which results from the co-moving frame transform.

The piecewise quadratic approximation of the spike profile leads to a piecewise constant curvature, which allows us to compute the perturbation exerted by an active axon on its neighbouring axons analytically. The curvature is given by the second derivative:

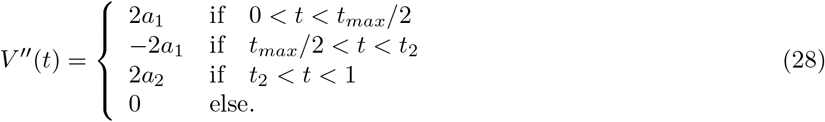

In the co-moving frame, this translates into

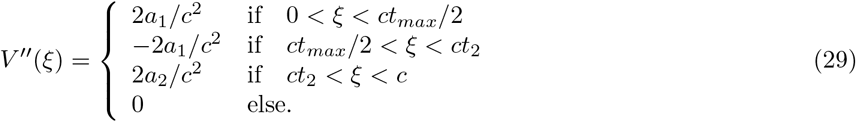

The quadratic approximation of the spike profile is shown in Figure 2, alongside its second derivative and the resulting perturbation profile in a passive axon. The perturbation profile can be computed by inserting Eq. (29) into Eq. (4), and inserting the resulting expression for 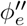 into Eq. (21).

**Figure 2:**
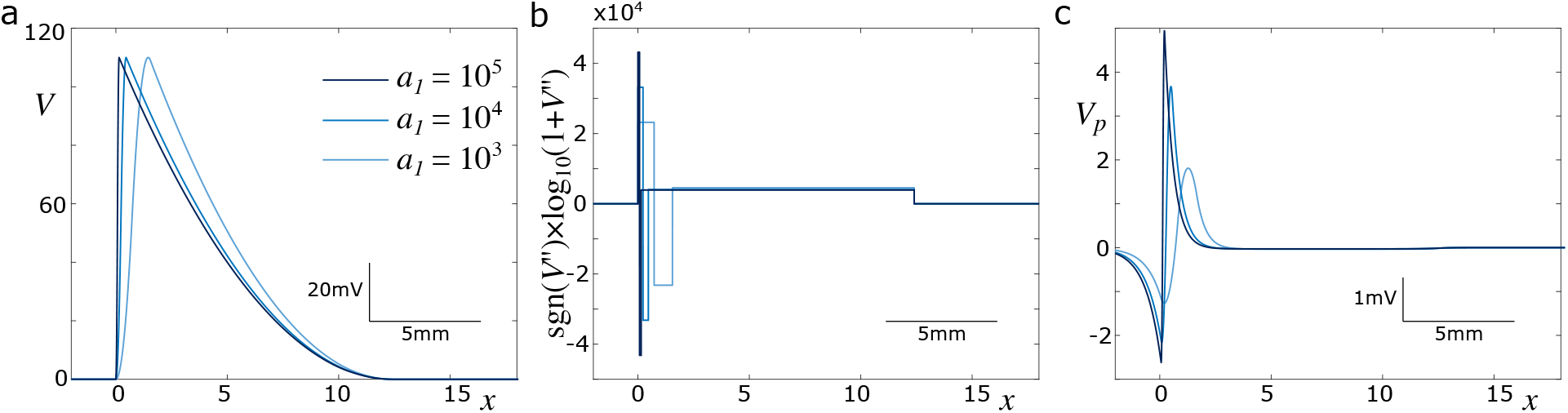
Piecewise quadratic approximation of spike profile. (a) The parameter *a*_1_ controls the shape of the spike profile, where larger values correspond to faster depolarisation. (b) The curvature (second spatial derivative) is piecewise constant. It is depicted on a signed logarithmic scale for better comparison. The unit of the curvature is mV/mm^2^. (c) Resulting perturbation in a passive axon. Parameters: *ρ* = 0.3, *d* = 1*µ*m, *v* = 3.1m/s, *T*_*s*_ = 4ms.

The complete mathematical expression for the perturbation in a passive axon driven by a spike in a second, active one is given by the following expression:

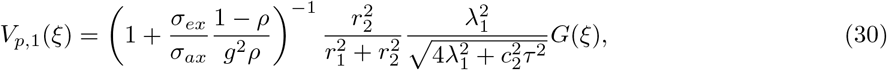

where *G*(*ξ*) is a function describing the spatial profile of the perturbation. The spatial profile can be formulated as follows:

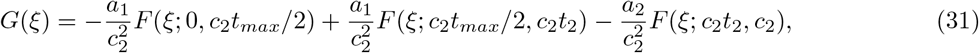

with

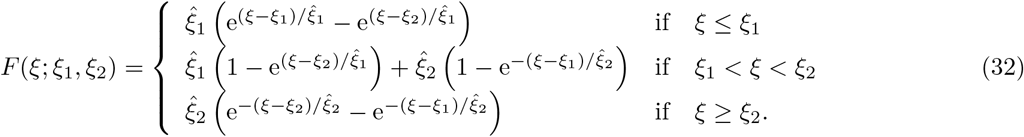

### 2.4 Calibration of spike propagation model

The spike propagation model was introduced previously for white-matter fibre bundles [26]. Here, we adapt this model to accommodate the specific properties of peripheral nerve bundles. The major difference lies in how the extracellular potential is generated. While in white-matter fibre bundles one has to consider their radial extent due to their large diameter, peripheral bundles can be regarded as quasi-one-dimensional objects in terms of the parameterisation of the extracellular potential. For instance, we consider bundles containing 100 axons with diameters of approximately 1*µ*m, which results in a bundle diameter of approximately 10*µ*m. The electrotonic length constant *λ* of such axons, however, is approximately 1mm. This means that the radial extent of a peripheral bundle can be neglected, because extracellular currents equalise radial charge imbalances almost instantaneously.

The core concept of the SPM is that a spike can be represented by its position on the axon, and its velocity. The velocity is considered constant along the axon in the unperturbed case. Perturbations of the membrane potential caused by spikes in other, contiguous axons, however, have an effect the propagation velocity, which is modelled in linearised form by:

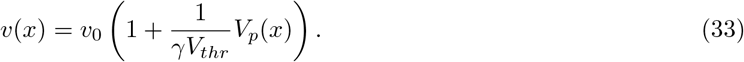

Here, the perturbation of the membrane potential, *V*_*p*_(*x*), is computed as shown in the previous section, which requires knowledge of the spike’s position. Finally, the position of the spike on the *i*^*th*^ axon (each axon carries at most one spike at any given time) is determined by

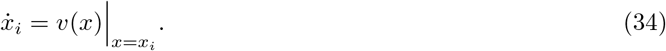

This means that the perturbation at the location of the spike determines the instantaneous propagation velocity. In general, Eq. (34) has to be solved numerically. For computational reasons, we consider the difference between spike positions to compute *v*(*x*):

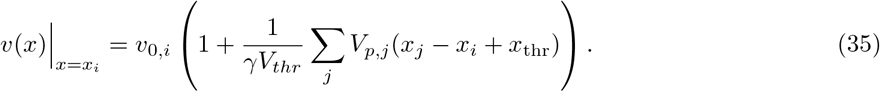

Since the spike position *x*_*i*_ refers to the point where the membrane potential first deviates from zero, 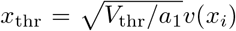 is introduced to mark the position where the membrane potential first reaches *V*_thr_. Consequently, the parameters *γ* and *V*_thr_ cannot be lumped together and have to be treated separately in the calibration.

In total, the SPM has three free parameters, which are the shape parameter *a*_1_, and the spike threshold *V*_*thr*_ and the coupling parameter *γ*, which set how strongly the perturbation affects the propagation velocity. The spike threshold also defines the spike position along the spike profile, where the spike is sensitive to perturbations. (We note here that although there are two threshold crossings, we only consider the one along the rising phase, which determines the propagation velocity. The falling phase (‘tail’) of the spike is due to repolarisation processes, which do not affect the spike velocity.)

To identify realistic values for the free parameters, we first generate data for the propagation velocities in two coupled axons using a biophysically realistic model, which is explained in detail in the next section. We use different parameter values of the fibre density and the fibre diameters. Specifically, the fibre density is chosen at *ρ* ∈ {0.2 0.4 0.6}, and the fibre diameters are chosen to be 1*µ*m for the smaller axon, and (1 + Δ*d*)*µ*m for the larger axon, with Δ*d* ∈ {0.1 0.3 0.5}. Thus, we fit the SPM to all nine parameter combinations 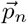.

To identify the best-fitting set of parameters, we define the following cost function that is to be minimised:

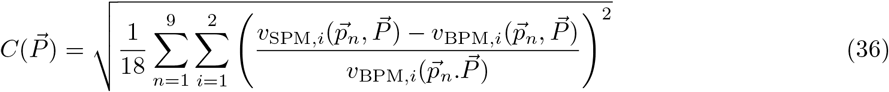

This cost function characterises the model discrepancy for a given set of free parameters 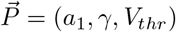.

In Figure 3 we illustrate the cost function for the three free parameters. There is a clear minimum at *a*_1_ = 800, which indicates that the optimal spike is relatively shallow. For the other two parameters we find a minimum at *γ* = 1.125 and *V*_*thr*_ = 19.5mV. This indicates that the perturbation of the membrane potential, *V*_*p*_, couples quite strongly into this expression. We note here that in our previous work [26], the optimal parameters were found to be *γ* = 6 and *V*_*thr*_ = 30mV. This discrepancy can be explained by the fact that there we used the extracellular potential *ϕ*_*e*_ to compute the perturbation, whereas here we use the perturbed membrane potential. The latter has a smaller amplitude due to axial currents, which is implicitly modelled by the convolution kernel.

**Figure 3:**
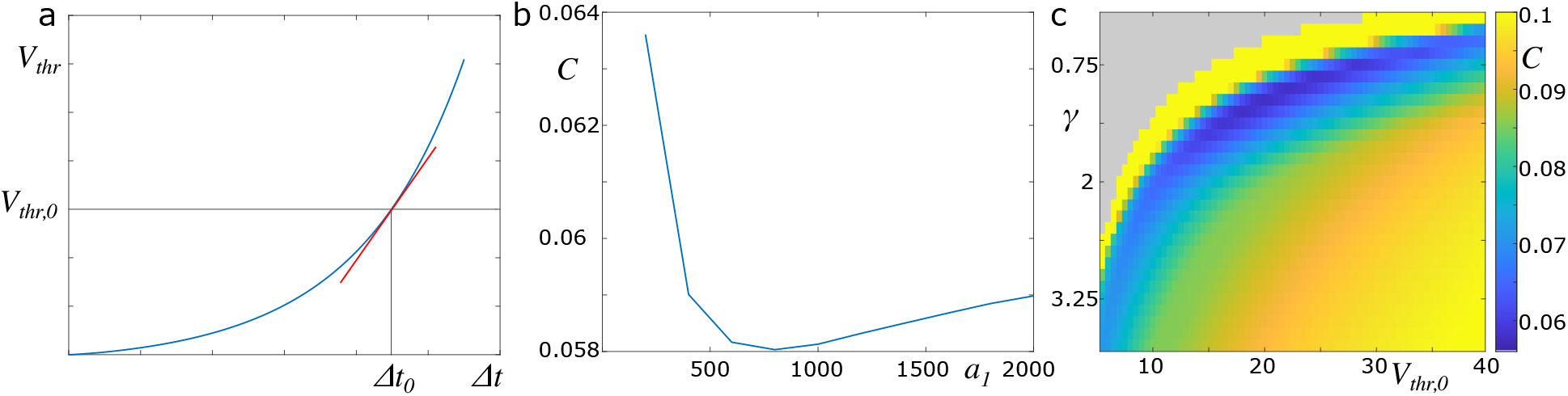
Calibration of spike propagation model. (a) An assumed nonlinear relationship between spike threshold and time lag between two reference points can be linearised around the default threshold value. The effect of perturbations of the membrane potential, and therefore of the spike threshold, on the time lag is then described in this linearised form. (b) Minimum of the cost function in *V*_*thr*_ and *γ* as a function of the shape parameter *a*_1_. The best fit occurs around *a*_1_ ≈ 800. (c) Cost function at *a*_1_ = 800 for varying values of *V*_*thr*_ and *γ*. The best fit is identified for the following set of parameters: *a*_1_ = 800, *γ* = 1.125, *V*_*thr*_ = 19.5.

### 2.5 Biophysical model of spike propagation

The biophysical model is an extension of Eq. (1), as it provides a detailed description of the nonlinear voltagegated currents. In this description, we distinguish between nodal and internodal segments. Internodal segments do not have voltage-gated ion channels, for which reason we set *I*_HH_(*V*) = 0. Voltage-gated ion channels are only located at the nodal segments, and they obey the following equations:

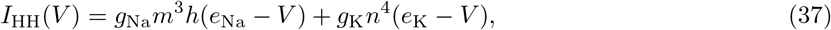

with the dynamics of the gating variables given by [30]

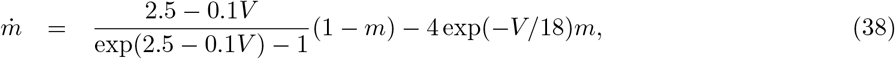

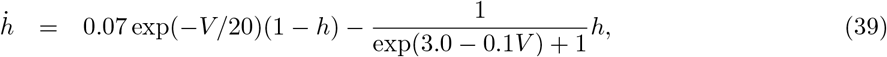

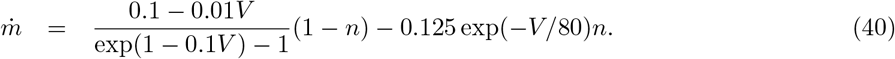

The reversal potentials in Eq. (37) are chosen to be *e*_Na_ = 115mV, and *e*_K_ = −12mV [31]. While the maximum conductivities were set to *g*_Na_ = 1200mS/cm^2^ and *g*_K_ = 90mS/cm^2^ in [31], we chose *g*_Na_ = 4800mS/cm^2^ and *g*_K_ = 720mS/cm^2^ to facilitate faster spike propagation.

## 3 Ephaptic coupling in nerve bundles

In this section we utilise the SPM to study the effect of fibre heterogeneity on the velocity of, and synchronisation between spikes. A special focus here is on fibre bundles with heterogeneous fibre diameters, which are distributed according to either a shifted, uniform distribution or a shifted alpha-distribution. Of particular interest is the interplay between this heterogeneity and the strength of ephaptic coupling as expressed by the fibre density.

### 3.1 Coupling between multiple nerve fibres

The expression for the effect of a spike carried by one axon on a neighbouring axon (Eq. (21) and Eq. (33)) can be extended to the perturbation in axons in larger axon bundles caused by multiple spikes. The perturbation of the membrane potential of the *i*^*th*^ axon is cumulative due to the assumed linearity:

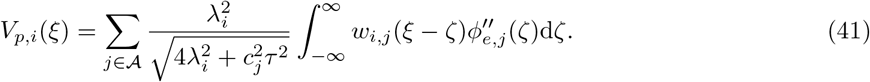

Here, 𝒜 denotes the set of active, spike-carrying axons, which is a subset of all axons within the bundle. The expression for the perturbation of the velocity in the *i*^*th*^ axon is then given by:

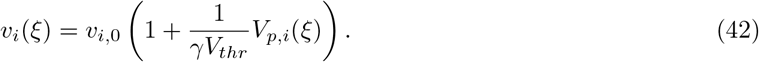

For simplicity we focus here on the case of spike volleys that engage all axons in the fibre bundle (i.e. 𝒜 is identical to the set of all axons), and that are completely synchronous initially (i.e. emission times are identical). The novelty here lies in the fact that the SPM reduces the computational effort as compared to the biophysical representation of the axon bundle, which allows us to model the interaction within spike volleys in large bundles. The numerical efficiency allows us to perform parameter screenings to describe the behaviour of the SPM in detail. Specifically, we vary the relative amount of extracellular space, which scales inversely with the amount of ephaptic coupling; and we vary the width of the distribution of axon diameters. We compare two types of distributions: a uniform distribution, and a shifted alpha distribution.

### 3.2 Uniform distribution of fibre diameters

The uniform distribution is set up as follows:

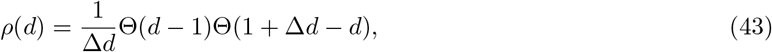

where Δ*d* is the width of the uniform distribution, and Θ(·) is the Heaviside step function. The minimum diameter is fixed at 1*µ*m, which results in a maximum diameter of (1+ Δ*d*)*µ*m in this distribution. The mean and standard deviation of this distribution is 1 + Δ*d*/2 and 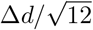, respectively. The coefficient of variation (standard deviation divided by mean) is thus 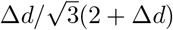.

In Figure 4 (a,b) we show the effect of fibre density and diameter distribution on the mean and standard deviation of the axonal delays. At small values of Δ*d* and large values of *ρ* spikes synchronise completely, as indicated by a standard deviation of delays close to zero. This is concurrent with an increase of the mean delay as the fibre density increases. The same synchronisation can be observed at larger values of Δ*d*, yet a higher fibre density *ρ* is required to achieve full synchronisation. Here, before full synchronisation sets in, the standard deviation of delays increases with *ρ* across a wide range, which indicates that there is no (complete) synchronisation of the spikes within the volley. Rather, the spikes tend to organise in clusters, which are synchronised internally. We show a delay density plot in Figure 4 (c), which illustrates the delay distribution across *ρ* for Δ*d* = 0.3*µ*m. Here, complete synchronisation occurs at *ρ* ≈ 0.9, yet already at *ρ* ≈ 0.6 one can observe clustering and slowing down of spikes. The participation of spikes in the different clusters is further illustrated in Figure 4 (d), which shows that spikes on axons with similar diameter tend to be part of the same cluster.

**Figure 4:**
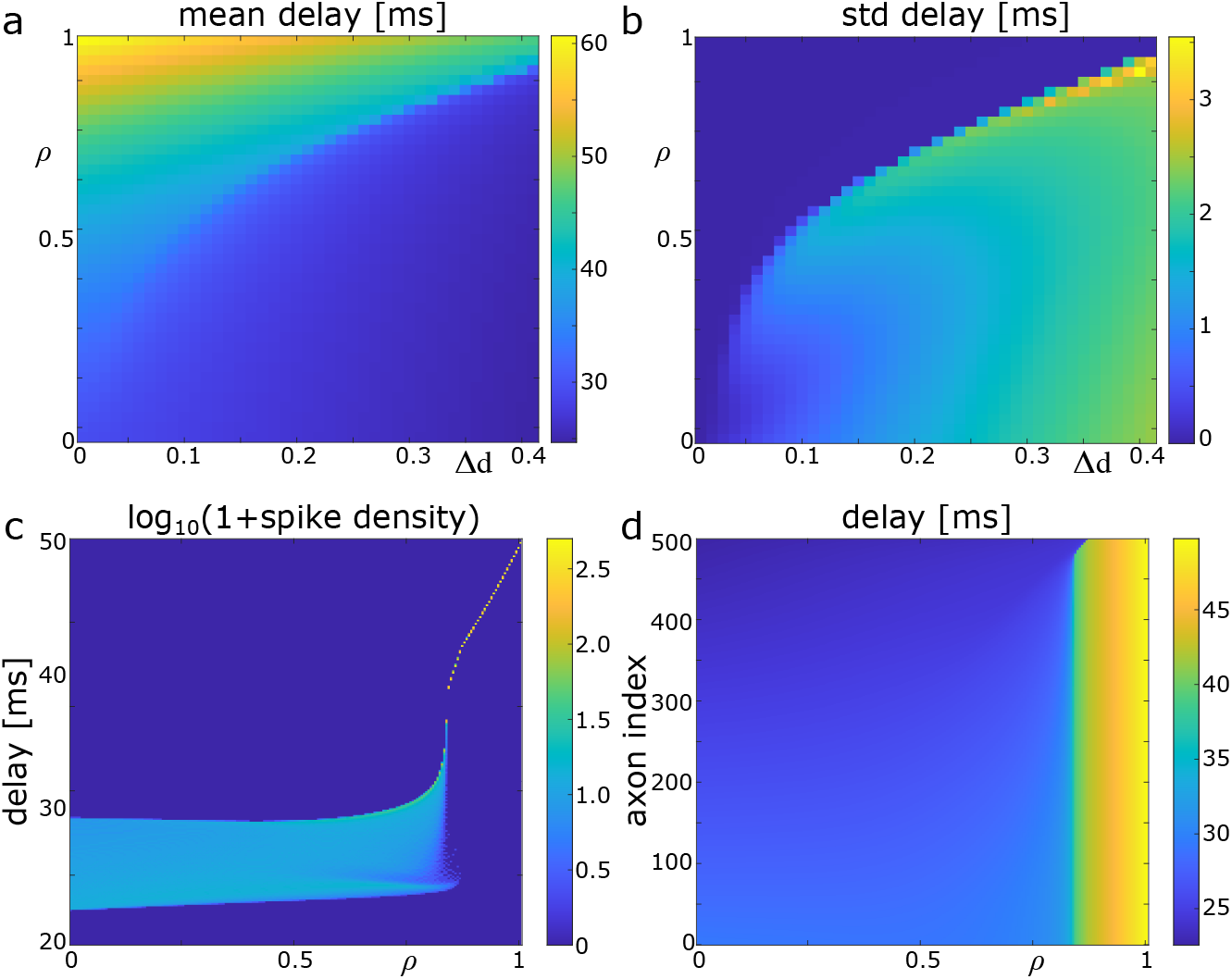
Delays in an axon bundle with uniform distribution of axonal diameters. (a) Mean axonal delay for varying values of Δ*d* and *ρ, N* = 100. (b) Standard deviation of delays, for same parameter range as in (a). (c) Logarithmic spike density showing the distribution of delays as *ρ* increases. Δ*d* = 0.3, *N* = 500. (d) same as (c), but with delays plotted across axon indices.

### 3.3 Fibre diameters from a shifted alpha-distribution

To test whether these results depend on the type of diameter distribution, we perform the same analysis for a shifted alpha distribution:

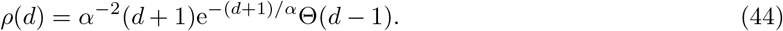

The minimum value in this distribution is *d* = 1, but the maximum diameter depends on the number of axons *N*.

Once more, we find an parameter regime at small values of *α* and large values of *ρ* where the spikes completely synchronise (Figure 5(a,b)). At larger values of *α* we find again that synchronisation is only partial, in line with the results obtained for the uniform distribution of diameters. This indicates that the specific type of distribution is not important, rather the width of the distribution is essential for the types of synchronisation (complete or partial) between the spikes. To illustrate the route to synchronisation, we detail the results for *α* = 0.03 in Figure 5(c). Once more, the spike density plot shows that across a wide range of *ρ*, synchronisation is partial, and only at large values of *ρ* it becomes complete. Differently from the uniform distribution, there is no clear clustering visible. The main cluster is formed by the spikes on the smallest axons. Figure 5 (d) shows as the fibre density increases, first the spikes in small axons synchronise, until all spikes are synchronous.

**Figure 5:**
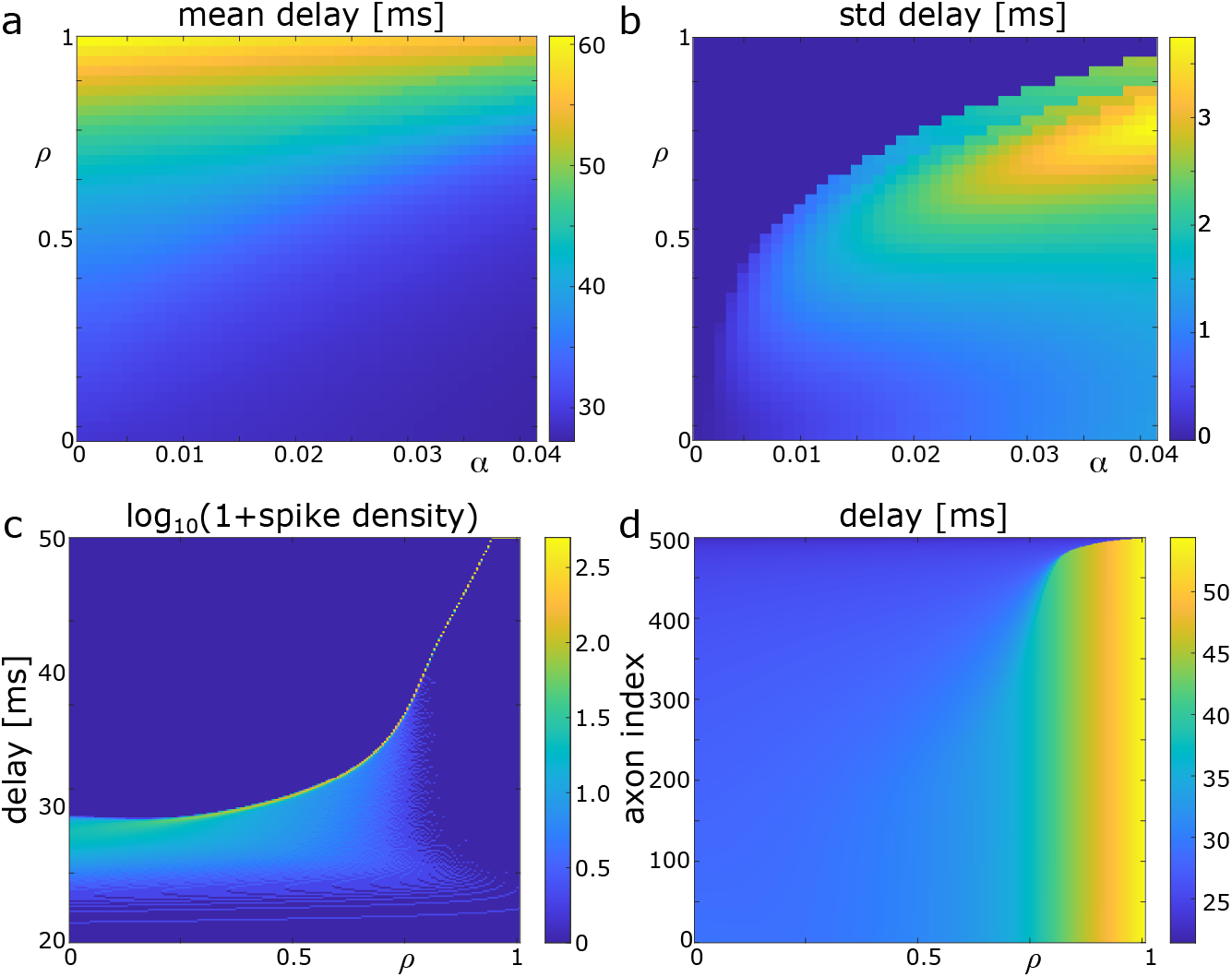
Delays in an axon bundle with axonal diameters obeying a shifted alpha distribution. (a) Mean axonal delay for varying values of *α* and *ρ, N* = 100. (b) Standard deviation of delays, for same parameter range as in (a). (c) Logarithmic spike density showing the distribution of delays as *ρ* increases. *α* = 0.03, *N* = 500. (d) same as (c), but with delays plotted across axon indices.

## 4 Discussion

The main contribution of the present study is to illustrate the effect of ephaptic coupling and fibre heterogeneity on the synchronisation of spike volleys in peripheral nerve bundles. The results confirm our initial hypothesis that while ephaptic coupling facilitates synchronisation, fibre heterogeneity counteracts this tendency. This is illustrated by the finding that for increasing fibre heterogeneity, the strength of ephaptic coupling (which increases with fibre density) necessary to produce full synchronisation within a spike volley also increases. This effect is independent of the specific type of fibre distribution chosen. At sufficiently high levels of heterogeneity, spike volleys no longer synchronise, even if the nerve bundle is completely filled with nerve fibre (fibre density *ρ* = 1). Nevertheless, in the absence of full synchronisation a more subtle phenomenon can be observed, which is the clustering of spikes into faster and slower spike packets. This is more prominent for the uniform fibre diameter distribution than the shifted alpha distribution, presumably due to the lack of a peak in the uniform distribution around which spikes could more easily cluster.

To obtain these results for large numbers of axons, we have adapted a spike propagation model (SPM) that was previously devised to simulate spike volleys in white matter fibre bundles [26]. The core assumption of the SPM is that the propagation of a spike is fully characterised by its intrinsic velocity (determined by structural and electrophysiological parameters of the corresponding nerve fibre), and the extracellular potential generated by spikes in nearby nerve fibres. The SPM therefore represents a much simpler model, without having to solve the nonlinear partial differential equation associated with the full biophysical model. Nonetheless, the SPM contains three free parameters that had to be calibrated using the biophysical model. We chose the scenario of two ephaptically coupled nerve fibres to generate data for delays with the biophysical model, and the discrepancy between the data and the output of the SPM defined a cost function. Minimising this cost function, we were able to find an unambiguous optimal set of parameters for which the SPM matched best the biophysical results. Regardless of the specific choice of parameters, we surmise that synchronisation and clustering of spike volleys is a universal phenomenon in the SPM.

Some limitations apply to the SPM. One main limitation is that spike volleys travel across the fibre bundle without the possibility of generating (ectopic) spikes, or extinguishing them. In the present study, we only consider spike volleys that engage all axons, therefore only the latter would be a possibility. A mechanism of spike generation / extinction could be incorporated into the SPM by defining conditions involving the perturbation of the membrane potential. More precisely, if the membrane of a passive axon is sufficiently depolarised, then a spike is generated there; conversely, if the membrane is sufficiently hyperpolarised, a spike is extinguished. A previous study, based on a spatially extended FitzHugh-Nagumo model, has demonstrated the possibility of emerging ectopic spikes in ephaptically coupled axons [17].

Another limitation is that our study is based on a homogenised formulation of axonal morphology, which discounts the precise location and alignment of nodes of Ranvier. While a previous study has shown that the effect of ephaptic coupling was stronger when nodes are aligned, a staggered arrangement of nodes yielded the same qualitative results [12]. Node alignment could be taken into account in the SPM by generating data with the biophysical model for different levels of nodal alignment, and fitting the coupling parameter *γ* to these values. Alternatively, theoretical considerations as in Ref [12] could serve to modulate the ephaptic coupling strength accordingly.

Finally, we would like to contemplate the possibility to confirm our results experimentally. While it is difficult to record spikes directly in peripheral nerves, it is easy to record the LFP generated by nervous activity using surface electrodes. The LFP of a spike volley can be computed using the line-source approximation [28, 32]. A typical LFP waveform generated by a single spike is triphasic, with an initial hyperpolarisation, intermediate depolarisation, and final hyperpolarisation. In the case of spike volleys, this wave form is convolved by the spatial distribution of spikes in the volley. As a consequence, highly synchronised volleys will show a strong triphasic profile, whereas distributed volleys show weaker profiles. The clustering that we have observed would then likely result in a multiphasic LFP profile, where the number if minima and maxima is determined by the number of spike clusters. Interestingly, experiments show such multiphasic LFP profiles, which hints at the existence of such a clustering regime [33]. Nevertheless, it is also possible that the multiphasic nature of these LFPs is due to feedback mechanisms, and it requires more detailed modelling work to disentangle the different mechanisms.

## Acknowledgements

HS was supported by a German Research Council (DFG) grant (No. KN 588/7-1, awarded to TRK), within priority program Computational Connectomics (SPP 2041).

